# Quantitative Fluorescence Imaging of Porphyrin Phospholipid Photobleaching and Light Activated Liposomal Doxorubicin Release Using Wide-field and Laparoscopic SFDI in an Ovarian Cancer Model

**DOI:** 10.1101/2025.02.14.638180

**Authors:** Rasel Ahmmed, Elias Kluiszo, Semra Aygun-Sunar, Matthew Willadsen, Hilliard L Kutscher, Jonathan F. Lovell, Ulas Sunar

## Abstract

Chemophototherapy (CPT) is an emerging cancer treatment that leverages the synergistic effects of photodynamic therapy (PDT) and chemotherapy. This approach utilizes photosensitizers like Porphyrin Phospholipid (PoP) and Doxorubicin (Dox) to enable phototriggered drug release and targeted tumor destruction. In this study, we present the development and validation of a wide-field laparoscopic spatial frequency domain imaging (SFDI) system, designed to improve intraoperative quantitative fluorescence imaging and monitoring of PoP photobleaching, a PDT-driven effect for tumor destruction, and light-activated Dox release, which facilitates targeted chemotherapeutic drug delivery in an ovarian cancer model. Compared to previous flexible endoscopic imaging methods, our laparoscopic SFDI system offers enhanced spatial coverage, enabling accurate wide-field optical property quantification in minimally invasive surgical settings. Using this system, we performed quantitative fluorescence imaging in vivo to obtain absolute concentrations of PoP and Dox fluorescence, correcting for tissue absorption and scattering effects. This capability allows for precise assessment of PoP photobleaching and Dox release kinetics with improved spatial resolution. Fluorescence imaging revealed a significant reduction in PoP concentration in tumor regions post-illumination, demonstrating the PDT-mediated photobleaching effect and successful light-triggered drug release activation for chemo-induced tumor destruction. The ability to differentiate PoP and Dox fluorescence in a laparoscopic system underscores its potential for real-time intraoperative monitoring of CPT efficacy. These findings establish wide-field laparoscopic SFDI as a promising tool for guiding minimally invasive photodynamic therapy and targeted drug delivery in clinical settings.

## 1. Introduction

Ovarian cancer remains one of the most lethal gynecological malignancies due to its high recurrence rates and peritoneal micrometastases, which are challenging to detect and treat effectively. Even after surgical treatment and adjuvant chemotherapy, up to 60% of patients may still exhibit occult disseminated ovarian disease[1-3], and systemic chemotherapy often leads to significant toxic side effects [4-7]. In addition, traditional imaging techniques, such as computed tomography (CT), magnetic resonance imaging (MRI), positron emission tomography (PET), and ultrasound, demonstrate less sensitive detection than reassessment surgeries [3, 8]. Thus, an unmet need exists for both sensitive imaging technologies for improved detection and therapeutic approaches that selectively treat superficial tumors in the intraperitoneal cavity while avoiding healthy tissue.

Photodynamic therapy (PDT) has gained significant attention as a minimally invasive cancer treatment modality that leverages light, a photosensitizing agent, and molecular oxygen to generate cytotoxic reactive oxygen species (ROS) capable of selectively destroying tumor cells [9] . Porphyrin-based compounds (photosensitizers-PS) are used for PDT and imaging because they strongly fluoresce, preferentially accumulate∼2-3 fold in malignant cells, and have demonstrated some success in detecting subcutaneous tumors with qualitative imaging approaches [7, 10-14]. Porphyrin conjugate (Porphyrin-phospholipid-PoP) has been through over 10 human phototherapy clinical trials. Because superficial tumors can be on the surface and accessible via surgery, PDT is an attractive approach for fulfilling the adjunctive therapy role of surgery [15]. However, high normal tissue toxicity has been a major obstacle to the clinical translation of PDT for ovarian cancer. Seminal works by Hasan[6, 7, 16-19] and Mordon [1, 20, 21] groups have shown epidermal growth factor receptor (EGFR) and folate-targeted approaches reducing the peritoneal tissue toxicity of PDT by using semi-quantitative imaging approaches in animal models. Liposomal nanocarriers serve to improve the biodistribution and efficacy of a variety of cancer drugs, with the lipid envelope protecting drugs from degradation inside the body and allowing for passive diffusion into tumor sites via the enhanced permeability and retention (EPR) effect[13, 22, 23]. While liposomal formulations can decrease side-effects, drug delivery to discrete tumor sites is suboptimal from intravenous administration alone [24]. The light activated PoP-liposomes used [25, 26] improve the concentration of drug delivered to the tumor site by selectively permeabilizing the lipid envelope with NIR light applied [27-30]. Both the lipid envelope of this drug as well as doxorubicin itself are inherently fluorescent, so fluorescent imaging of the drug before and after light activation can quantify drug kinetics in vivo [31].

To develop an image-guided system for the delivery and detection of PS, knowledge about PS concentrations at and near the target cells is essential. Our approach capitalizes on the high sensitivity and specificity of fluorescence contrast, the fact that Porphyrin is highly fluorescent [32], and the increased resolution of wide-field imaging. However, the accurate quantification of drug concentrations in tumors remains an ongoing challenge because the raw fluorescence signal is affected by tumor absorption and scattering properties, which confound the true fluorescence contrast [33, 34]. To correct this distortion, spectroscopic measurements can be taken to quantify tissue attenuation, but it is point-specific [35-38]. SFDI gathers tissue data over a wide area, allowing for accurate measurements in heterogeneous tissue like the intraperitoneal cavity.

SFDI can quantify both optical absorption and scattering during reflectance imaging mode [7] and easy to implement [39]. Knowledge of the optical parameters can allow modeling of the light dose distribution within the treatment field. In addition to light dosimetry, these parameters can provide the fluorescence correction factor for tissue attenuation so that one can extract absolute fluorescence concentration in vivo.

We employed a custom laparoscopic SFDI system to quantify the absolute concentration of PoP and Dox fluorescence concentrations in an ovarian cancer model. Doxorubicin (Dox), embedded in the liposomal construct of PoP enables spatially and temporally controlled release of Dox from liposomes using near-infrared light [40]. The quantification of PoP fluorescence concentration revealed a significant photobleaching effect compared to the periphery. Then, we monitored the release of Doxorubicin (Dox) and demonstrated significant release at the target sites compared to surrounding tissue.

## 2. Results

### 2.1. PoP Fluorescence Concentration Contrast and Photobleaching in Subcutaneous Tumors

Figure 1 provides a comprehensive visualization of the optical characteristics of the tumor and its surrounding periphery. Figure 1(a) and 1(b) show the diffuse reflectance image with the lesion and surrounding periphery area. These images offer insight into the spatial variations in light scattering and absorption within the tissue. Whereas 1(c) and 1(d) show the absorption and scattering maps at 660 nm, respectively. Fig. 1(c) showed higher absorption at the lesion compared to the surrounding periphery likely due to enhanced vascularization or chromophore concentration in the tumor. Conversely the scattering parameter at 660 nm was lower at the tumor in Fig. 1(d) suggesting structural alterations such as reduced collagen density or cellular disorganization. Only one wavelength (660 nm) is presented for clarity; the 660 nm wavelength was chosen because this is the excitation wavelength of PoP used for PDT and Dox release.

**Figure 1.**
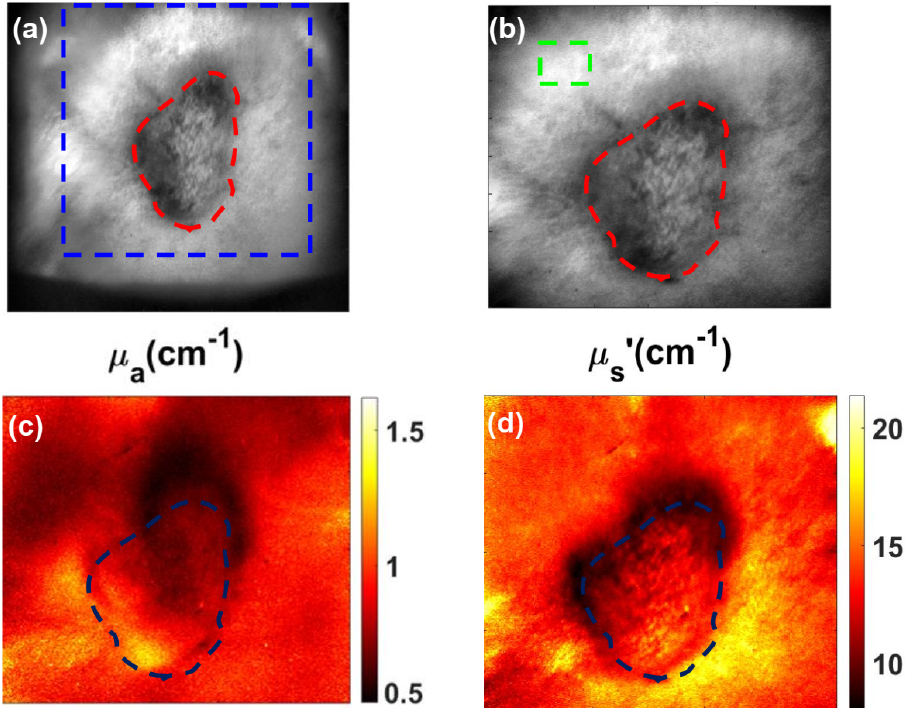
Representation of optical properties of the tumor and periphery (**a**) Raw image with a specific region of interest (ROI) including both periphery and tumor region. (**b**) Diffuse reflectance image with the lesion and periphery marked. (**c** and **d**) Absorption and scattering map for 660 nm.

Figure 2 shows representative images of a tumor in a mouse 1h after administration of PoP. Figure 2(a) shows the white light image to indicate the tumor location. The tumors were large (∼9 mm diameter). The uncorrected fluorescence shows unexpected tumor contrast (705.09±94.2a.u.-tumor vs 983.07± 58.51a.u.-periphery). This suggests that raw fluorescence intensity alone does not accurately reflect PoP distribution due to light attenuation in tissue. However, after applying a correction algorithm to quantify fluorescence, the quantified absolute PoP concentration image demonstrated a higher contrast in the tumor, compared to peripheral tissue (0.24±0.026 µg/mL vs 0.183±0.0001 µg/mL, respectively). This corrected fluoresnece concentration confirms that PoP preferentially accumulates in the tumor, enhancing tumor contrast and supporting its potential role in targeted photodynamic therapy (PDT) and drug delivery applications.

**Figure 2.**
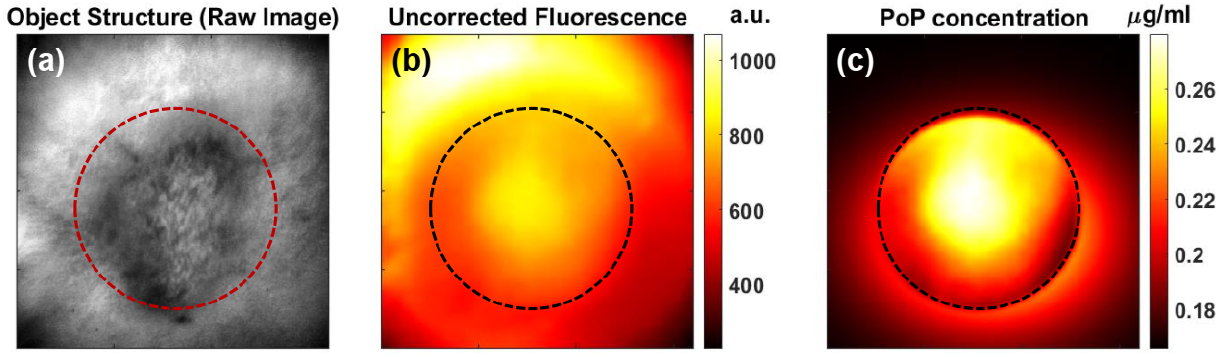
Representative images of a tumor after light administration to PoP (**a**) White light structural image showing the tumor area. (**b**) Uncorrected fluorescence image does not show localized contrast. (**c**) PoP fluorescence concentration indicated higher contrast between the tumor and the surrounding area compared to the uncorrected fluorescence.

We also quantified the PS photobleaching in vivo. As shown in Figure 3(a), the PoP fluorescence amplitude was approximately 2.2 times higher in tumor tissue than in normal tissue before PDT. Following treatment light irradiation, fluorescence imaging revealed a significant (∼75%) reduction in PoP concentration in the tumor region due to photobleaching [41-46]. Figure 3b shows the absolute PoP concentration, indicating greater PoP uptake in tumor tissue compared to normal tissue, with a tumor-to-normal tissue uptake ratio of approximately 1.7, consistent with previously reported values[8] . After PDT, the PoP concentration in the tumor decreased by about ∼25% due to photobleaching after treatment, while the drug concentration in normal tissue remained similar to the pre-PDT value.

**Figure 3.**
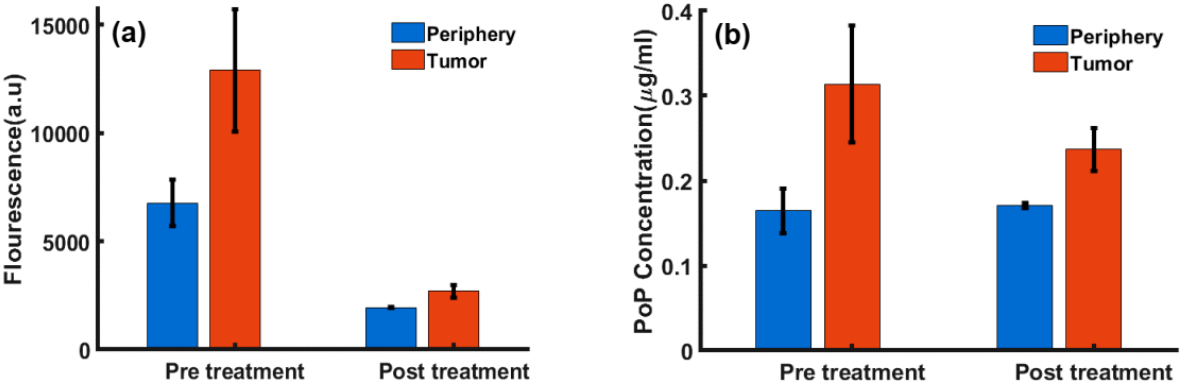
Extracted functional parameters from tumor and periphery before and after PDT (**a**) Bar plot showing the comparison of PoP fluorescence in the tumor vs peripheral tissue pre and post-treatment light (**b**) Comparison of the PoP concentration (µg/ml) before and after treatment.

### 2.2. Dox release in Mouse Carcass

#### 2.2.1. Dox imaging contrast

We then investigated Dox imaging contrast in a BALB/c mouse carcass. Figure 4(a) highlights the white light structural image, marking the PoP injection and Dox release sites. The injection site was exposed to treatment light for 8 minutes. The uncorrected fluorescence in Figure 4(b) and concentration map in Figure 4(c) revealed differences in image contrast. This is due to attenuation correction in the tissue’s optical properties differences between the skin vs Intralipid. Additionally, autofluorescence was subtracted from the total fluorescence signal to isolate the Dox contribution. Autofluorescence accounted for approximately 60% of the initial background signal before treatment and about 10% of the peak fluorescence signal during treatment.

**Figure 4.**
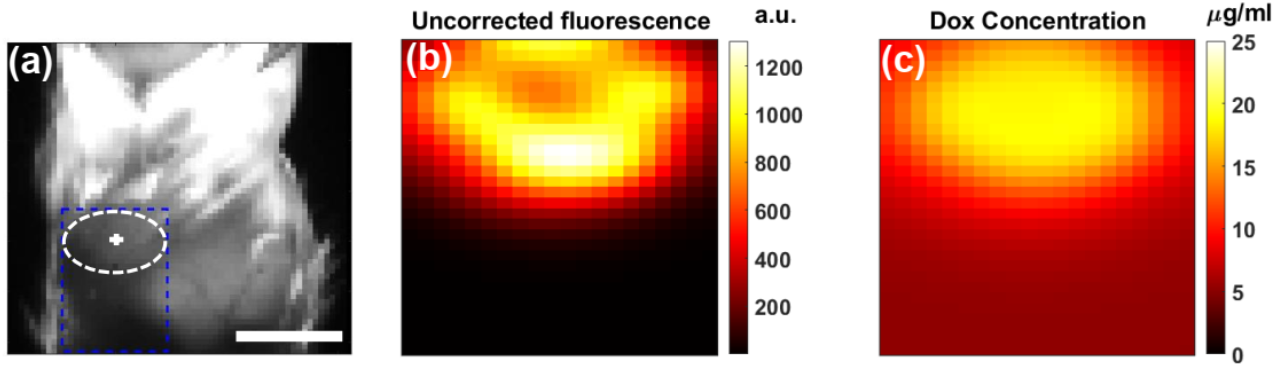
Representative Dox quantification in a mouse carcass. (**a**) White light image showing the structure of the PoP injection and release site. (**b**) Uncorrected Dox fluorescence image after 8-minute illumination. (**c**) Dox concentration image after 8-minute illumination

##### 2.2.2. Dox release kinetics

We examined Dox release kinetics in a mouse carcass Figure 5. Figures 5(a), 5(b), and 5(c) display Dox concentrations at pre-treatment, 4 minutes post-treatment, and 8 minutes post-treatment, respectively, showing an increase from 6.53 ± 0.74 µg/mL to 13.01 ± 1.24 µg/mL distribution, likely due to variations in optical parameters, particularly scattering differences between the skin and intralipid. This is likely attributed to the presence of overlying skin and the limited penetration depth of the excitation light. Additionally, it is expected that the treatment light did not fully probe the entire drug volume due to the partial volume effect caused by the skin layer and the uneven distribution of the pro-drug PoP beneath the skin. As a result, a portion of the total volume near the surface may have been released initially and subsequently diffused into the untreated regions over time.

**Figure 5.**
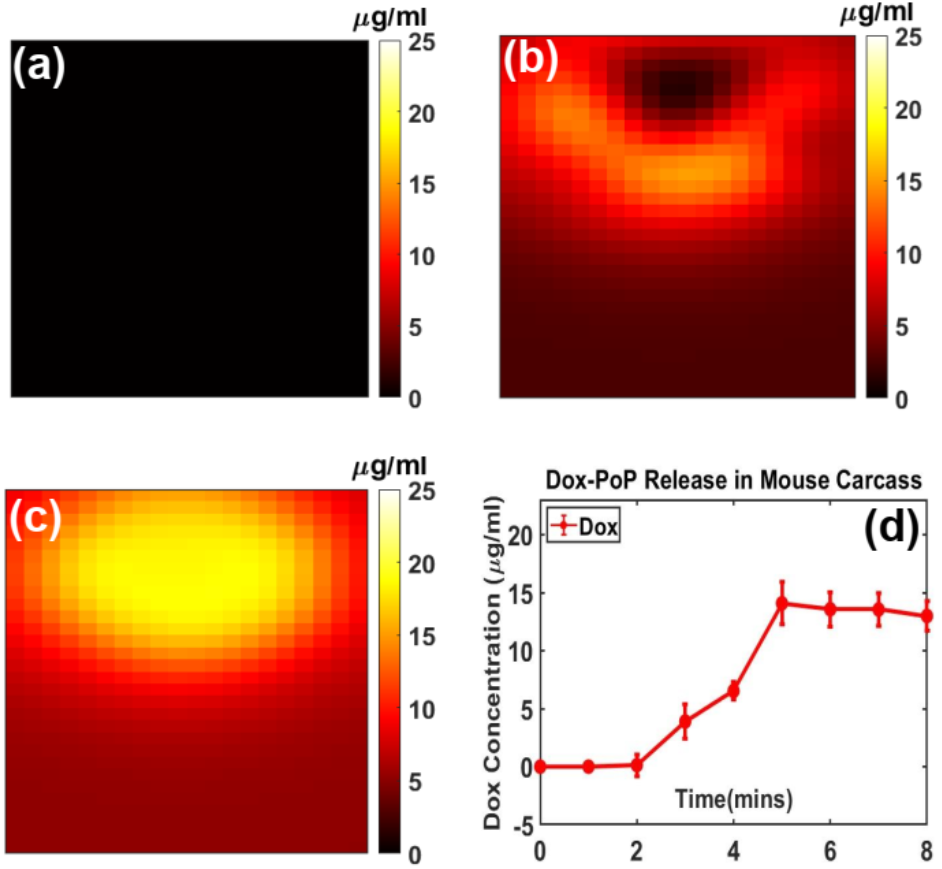
Dox releases kinetics in a mouse carcass. (**a**) Pre-treatment (**b**) 4 min post-treatment (**c**) 8 min post-treatment. (**d**) Complete Dox release kinetics curve with mean and standard deviation of the ROI.

##### 2.2.3. Porphyrin photobleaching in Dox-PoP drug

We compared Porphyrin fluorescence before (Figure 6 (a-c)) and after (Figure 6(d-f)) treatment light for Doxorubicin release. It showed a decrease from 1.57 ± 0.37 µg/mL to 0.73± 0.14 µg/mL, likely due to photobleaching from the strong treatment light at the excitation wavelength of the Porphyrin. These findings confirm that the treatment light not only facilitates Dox release but also impacts Porphyrin fluorescence (photobleaching), highlighting the importance of optimizing light dosimetry for controlled drug activation and optimizing photobleaching for the synergistic effect of chemotherapy and photodynamic therapy (PDT).

**Figure 6.**
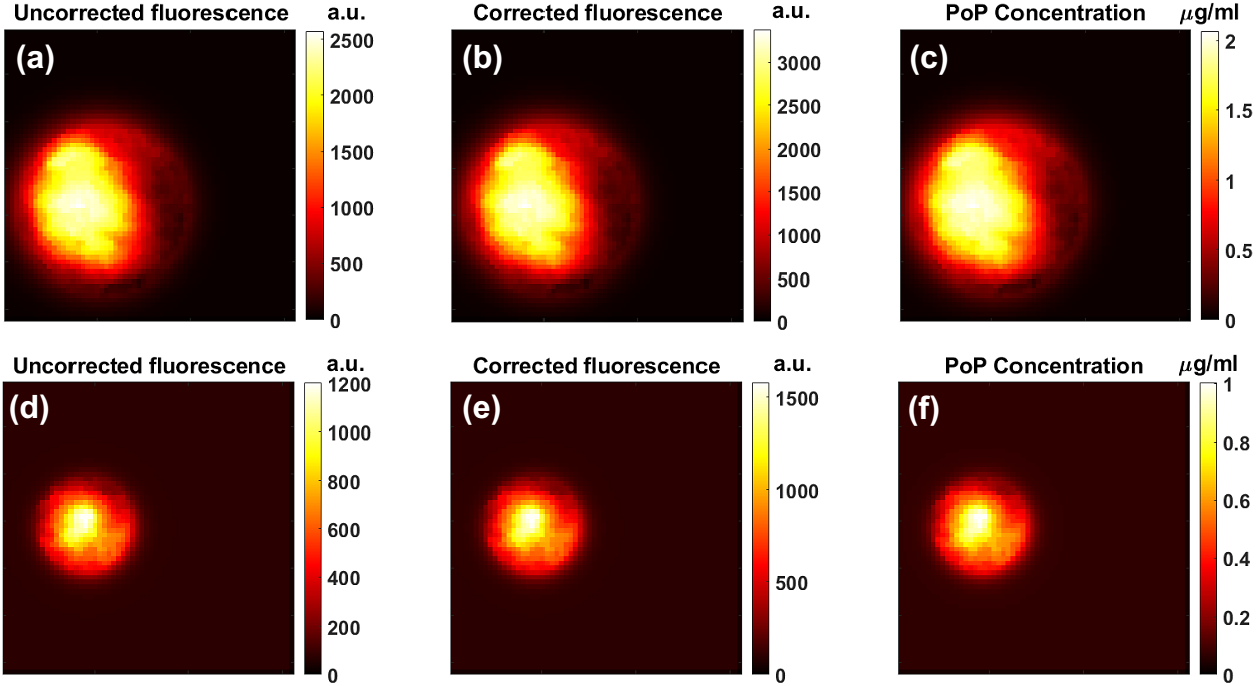
Porphyrin fluorescence pre- and post-light treatment. (**a**-**c**) pre-treatment fluorescence and subsequent PoP concentration, (**d**-**f**) post-treatment fluorescence and subsequent PoP concentration.

## 3. Discussion

In this study, our quantitative, wide-field laparoscopic approach was used in reflectance and fluorescence imaging modes to monitor subcutaneous administration of PoP and Dox photosensitizers. SFDI quantified the optical parameters of the model tissue to provide *a priori* information used in correcting fluorescence signal attenuation, leading to accurate models of drug concentrations. Quantitatively derived distributions of both PoP and Dox function as a guide for CPT treatment, with PoP photobleaching demonstrating the effects of PDT and Dox fluorescence showing the bioavailability of the chemotherapeutic.

However, achieving concurrent measurements during PDT remains complex. Unlike probe-based methods[26], which may interrupt treatment light and require fine adjustments during measurements, wide-field SFDI avoids physical interference with the treatment area. Furthermore, non-contact methods such as SFDI dual-channel endoscope systems [25] provide the flexibility to monitor tumors with flat surfaces, such as subcutaneous tumor models or skin, without physical disruption.

PoP fluorescence correction was crucial at the tumor site, where absorption and scattering parameters were ∼40% and ∼15% higher than the periphery, respectively (Fig. 2b and 2c). Thus, one could incorrectly conclude from raw fluorescence images alone that PoP distribution was less at the tumor site than the periphery, implying that in some cases, corrected fluorophore correction could be crucial in individual assessment of treatment efficacy.

The ∼660 nm treatment light activation was generated by a laser source and collimated through an SMA fiber to activate PoP at the treatment site. In areas with highly heterogeneous optical properties, can lead to significant variations in local light dose [47]. Thus, the knowledge of the tissue optical properties is important for both accurate drug dose distribution at pre-, during and post-treatment, as well as accurate local treatment light delivery and optimization on an individual, tumor-by-tumor basis. In future studies, the laser light source can be directed through the DMD. This could allow changes in the shape and intensity of the treatment light specific to each tumor, while still retaining the required fluence rate for Dox-PoP activation.

In the laparoscopic SFDI data, there was a leveling off of Doxorubicin fluorescence sthat occurred within 6 minutes of treatment time, likely due to the completed release of the PoP-liposomes. Notably, the total activated Dox concentration shown in Figure 5(d) did not reach the injected amount (equivalent to ∼23 µg/mL of Dox). This is likely due to the shallow penetration depth and partial volume effect issues in vivo measurements. The limited penetration depth of 490nm light might also explain the heterogeneity of the Doxorubicin signal in Fig. 5b, as liposomes closer to the surface of the injection could be released first and then gradually mixed with the untreated volume over a longer period.

In Figure 5, the targeted region was located at or near the surface, allowing for direct and efficient light delivery to achieve localized drug activation. However, in deeper anatomical sites, such as the peritoneal cavity, achieving uniform light exposure over a large area can be challenging. Alternative light delivery methods [48], including interstitial fiber-optic probes or endoscopic-based illumination [25], can provide viable solutions for enabling precise and effective light-triggered drug release in such scenarios. Overall, our SFDI system has shown a 10% error or less in calibration tests with tissue-mimicking phantoms. This error ultimately translates to the concentration maps of fluorophores, which require the absorption and scattering parameters at the excitation and emission wavelengths as inputs. To optimize SFDI measurements in vivo, future studies will incorporate profilometry techniques to account for the curvature of the tumor site. We did not correct for the surface angle variations since the injection and tumor size were relatively small, but the error is likely still present. For larger tumors, we would need to implement a model-based Lambertian reflectance approach for profile correction, as demonstrated by Gioux et al [49].

It should also be noted that all the reported values were obtained via post-processing with custom MATLAB software that fit the data to the model at each pixel. This process took between 5 to 10 minutes, depending on the pixel binning and was performed outside the clinic. In total, the data acquisition time was around 1 minute for SFDI, pixel-by-pixel fitting for optical properties took 1-2 minutes per wavelength, and the multi-wavelength fitting took 2-3 minutes. These times could be reduced by binning to fit fewer pixels. Utilizing a faster acquisition technique, such as a single snapshot [49] or a faster processing technique such as the lookup table (LUT)[49] model recently proposed by Angelo et al., [50] could reduce the overall quantification time and provide results in the clinic. This near-real-time feedback would be particularly useful for monitoring light-based therapies such as laser or photodynamic therapy.

## 4. Materials and Methods

### 4.1. Wide-field SFDI System

A clinic-friendly SFDI system was constructed as shown in Figure 7. The system utilized four high-power, compact light-emitting diodes (LEDs) from the LCS series, each emitting light at 590 nm, 625 nm, 660 nm, and 740 nm (Mightex, Toronto, Ontario, Canada). A four-channel LED controller (Mightex) sequentially activated the selected excitation wavelength. The light was directed via a liquid light guide to a projector (Light Commander; Logic PD, Inc., Minneapolis, MN, USA) equipped with a digital micromirror device (DMD) module offering a resolution of 1024 × 768 pixels.

**Figure 7.**
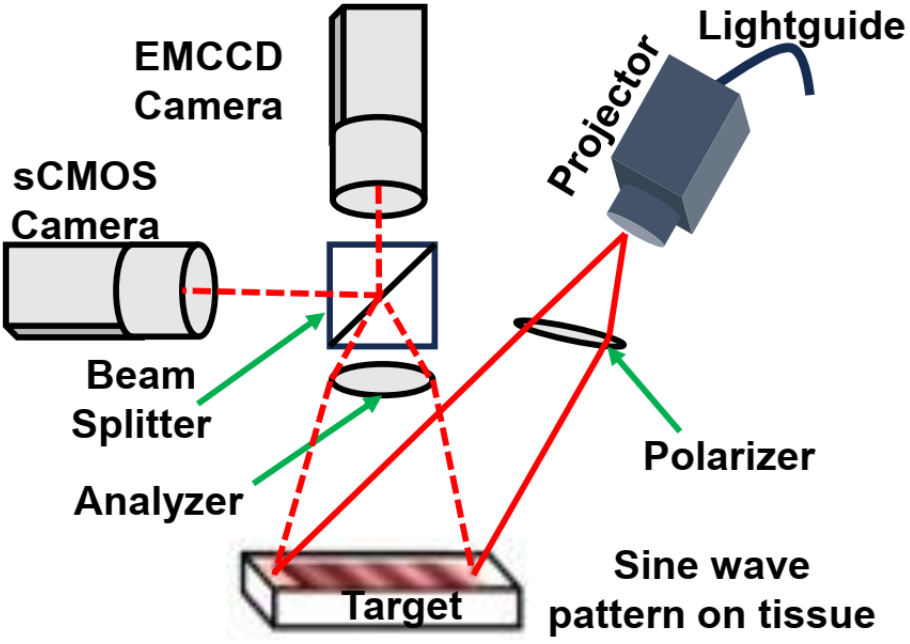
Schematic diagram of the imaging head showing the projector module, One charge-coupled device (CCD), and one sCMOS camera, beam splitter, polarizer, and analyzer. Light-emitting diode (LED) light is delivered with a light guide. Four LEDs are switched sequentially. Digital micromirror device generates sinusoidal patterns, patterns projected onto the skin surface by the projector, and the reflected signal is detected by CCD cameras

The DMD module generated sinusoidal patterns with three distinct phases (0, 2π/3, 4π/3) and 22 spatial frequencies ranging from 0 to 3.1764 cm^-1^. These patterns were projected onto the tumor surface, and the reflected light was captured by the cameras. The cameras were aligned to focus on the same field of view as the projector, covering an area of 22 × 22 mm^2^. A rigid light shield with an imaging window was used to block ambient light and maintain a consistent distance from the target tissue. For imaging, the system incorporated two cameras positioned on either side of a 685 nm dichroic mirror (67-085; Edmund Optics, Barrington, NJ, USA). This design ensured precise imaging of the illuminated area. Fluorescence and reflectance imaging were performed simultaneously. The first sCMOS camera (Zyla Andor, Belfast, Ireland) captured reflectance images at 590, 630, and 660 nm, while a highly sensitive Electron Multiplying Charge-Coupled Device (EMCCD) camera (Luca; Andor, Belfast, Ireland) acquired reflectance images at 660nm and 740 nm and fluorescence data at 740 nm. Splitting the light at the 685 nm dichroic mirror allows for analysis of the projected excitation light by the sCMOS, and the fluorescent signal by the more sensitive EMCCD camera.

The cameras operated with an acquisition time of 100 ms per image, resulting in a total acquisition time of 27 seconds (calculated as 100 ms × 3 phases × 22 frequencies × 4 wavelengths). The system was fully automated using a custom LabView (National Instruments, Austin, TX, USA) software, which included subprograms to manage all system components. The software enabled automatic adjustments of LED intensities and exposure times for individual subjects.

To minimize specular reflections during reflectance imaging, cross-polarizers were placed in front of the projector and cameras. The LED light source intensity was maintained at <1 mW/cm^2^ to ensure safety.

The optical absorption and scattering properties were quantified by fitting modified frequency-dependent Monte Carlo model to the measured reflectance data across multiple spatial frequencies. This process utilized a reference phantom with known optical properties, as previously described [51]. All 22 spatial frequencies, ranging from 0 to 3.1764 cm^-1^, were included in the analysis. For each spatial frequency and wavelength, the three phase-shifted reflectance images were demodulated to isolate the spatially modulated component of the diffuse reflectance. The demodulated reflectance, which varies as a function of spatial modulation frequency, exhibits differing sensitivities to absorption and scattering parameters depending on the frequency. This allows SFDI to independently and accurately quantify both absorption and scattering. Using this approach, pixel-by-pixel fitting was performed to generate spatial maps of absorption and scattering properties.

### 4.2. Laparoscopic SFDI System

The laparoscopic SFDI system was constructed as shown in Figure 8. Figure 8(a), showing the mouse carcass imaging setup, and Figure 8(b) detailing the projection and imaging components within the laparoscopic system.

**Figure 8.**
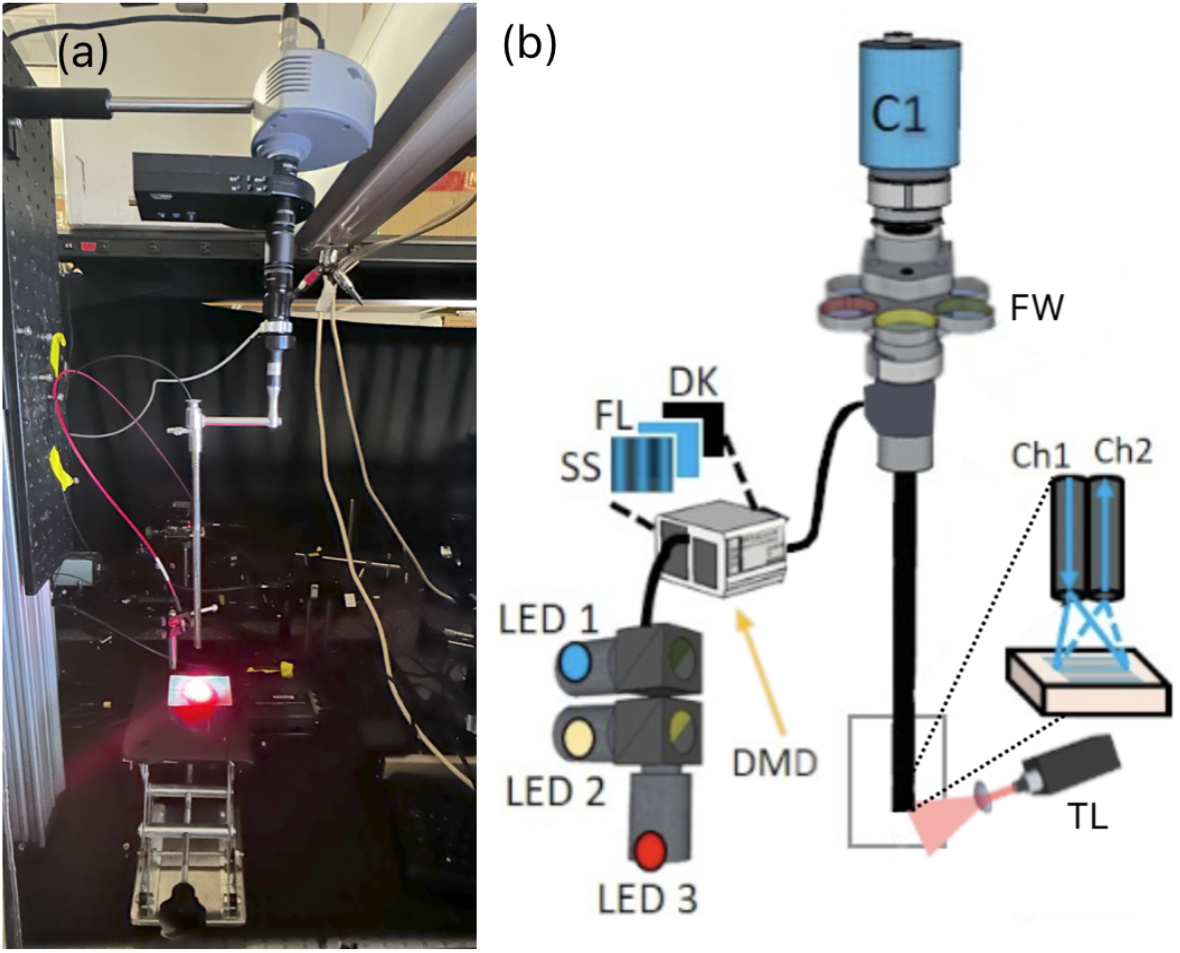
The experimental setup of Doxorubicin release in mouse carcass, (**a**) laparoscope projecting 490nm excitation light in blue and the 656nm treatment light in red, (**b**) laparoscope setup, showing LEDs and DMD with sinusoidal, fluorescent, and dark images projected, as well as the camera (C1), filter wheel placement (FW), treatment light (TL) and the projection (Ch1) and imaging (Ch2) channels.

Our laparoscopic fluorescence imaging and SFDI setup consisted of a modified DMD (LightCrafter 4500, Texas Instruments) to spatially modulate light to the appropriate sine wave patterns with three different phases (0, 2π /3, 4π/3) and five spatial frequencies from 0 to 2.0 cm^-1^ at a resolution of 1280× 800 pixels.

Light from three high-power LEDs at 490nm, 590nm, and 656nm was directed to the DMD through a liquid light guide (Mightex), with the LEDs and DMD being controlled remotely via MATLAB. The 490 nm LED excited Dox fluorescence and captured SFDI optical properties at its excitation peak, the 590 nm LED measured optical properties and Dox fluorescence at its emission peak, and the 656 nm light excited Porphyrin fluorescence.

The DMD module generated sinusoidal patterns and projected them through a 2.4 mm imaging fiber with 13,000 elements (Asahi Kasei, Tokyo, Japan) which passed through the fixed laparoscope to the distal end facing the imaged surface. A custom objective lens at the tip of the fiber served to collimate light from the fiber and homogeneously projected onto the tissue. Patterns from the DMD were reflected off the tissue and collected through the laparoscope optics, which includes the laparoscope (8912.43, R. Wolf), a zoom coupler (Accu-Beam, TTI Medical), two 30mm achromatic lenses, a filter wheel and an aperture (ThorLabs) before reaching the EMCCD camera (1004 ×1002 pixels, Luca, Andor, Belfast, Ireland). The camera was focused over the entire area of the projected SFDI pattern at a 3.2×3.2cm field of view (FOV). The optical design ensures an accurate sinusoidal projection for SFDI in a format using components currently used in laparoscopic surgery.

For acquiring SFDI data, no optical filters were used, with the LED output centered on the respective 490nm and 590nm excitation and emission wavelengths of Doxorubicin. When imaging Doxorubicin fluorescence, a 530nm long pass filter, and a 593±40nm fluorescent filter were used to isolate Doxorubicin fluorescence from the excitation light. When imaging Porphyrin fluorescence, a 660±10 nm bandpass filter was used to isolate the PoP excitation signal, and a 716±40nm fluorescence filter was used to separate PoP fluorescence. Fluorescent images were taken with a 6-seconds exposure time and total 12 seconds (including dark) at 16×16 camera binning. Whereas SFDI measurements were taken at 2 seconds exposure per projection, at 8×8 camera binning with a total time (3 phases × 5 frequency) of 30 seconds for each wavelength. For SFDI measurements cross-polarizers were built into the front of the laparoscope’s distal end to reduce spectral reflection. A structural frame acted as a mechanical stop to ensure that the object surface was no closer than the minimal effective working distance of the scope. Due to the divergence of light from the objective lenses, the frequency of the projected SFDI patterns will increase with distance, so the structural frame ensured that the target was always close to the optimal working distance. The optical absorption and scattering properties were extracted from SFDI images by modified frequency-domain Monte Carlo model to the measured reflectance data across multiple spatial frequencies as described prior but with a total of 15 images at 5 frequencies ranging from 0 to 2.0 cm^-1^.

### 4.3. System Calibration

The wide-field SFDI instrument was tested on ovarian tissue mimicking phantoms with optical absorption (µ_a_) and scattering (µ_s_’) properties within the range of tissue at 660 nm, which is the common wavelength for PoP-based PDT. The calibration phantoms used in this experiment were fabricated based on absorption and scattering parameters from prior work [52]. Bulk optical parameters were quantified by fitting frequency-dependent reflectance data with modified frequency-domain diffusion models by using a reference phantom with known optical properties [51].

The laparoscopic SFDI system was characterized with multiple calibration phantoms made by titrating the optical properties ranging from µ_a_= 0.5cm^-1^to 1.5cm^-1^and µ_s_’ = 10cm^-1^to 30cm^-1^at 490nm and 590nm. The mean percent error in quantifying the absorption parameter was always less than 10%. Fluorescence phantoms were prepared by titrating the Dox concentration, with a 0.5mg/mL free-Dox solution used to acquire fluorescence values at 2, 4, 6, and 8 µg/mL in phantoms of varied optical properties. Reconstructed raw fluorescence values for concentration were used as a calibration curve to obtain absolute Dox concentrations as previously described [53].

### 4.4. Subcutaneous tumor culture in mouse model

Human epithelial ovarian adenocarcinoma cell lines SKOV3 were purchased from the American Type Culture Collection (ATCC, HTB-77, Manassas, VA, USA). The cells were maintained in McCoy’s 5A medium supplemented with 10% Fetal Bovine Serum and 1% antibiotic-antimycotic. Cells were cultured at 37 °C in a humidified atmosphere of 95 % air and 5 % CO_2_ (v/v). Cells were routinely sub-cultured when 90% confluence was reached using 0.25% w/v Trypsin-EDTA solution. Cell culture medium and supplements were purchased from Gibco-Life Technologies (Grand Island, NY, USA)

*In vivo* studies were conducted by an Institutional Animal Care and Use Committee (IACUC) of Wright State University and Stony Brook University-approved protocols. All animal care was performed in accordance with the relevant guidelines and regulations outlined in the “Guide for Care and Use of Laboratory Animals.

Female athymic *nu*/*nu* (nude) mice were purchased from Charles River Laboratories. When they were 8-10 weeks old, they were injected subcutaneously (*sc*) with 10^7^SKOV3 cells. Mice were then monitored for the growth of tumors, and tumor diameters (length; L and width; W) were measured with a caliper, Tumor volume was calculated using the following formula: (L×W^2^)/2. When tumors reached approximately 9 mm in diameter, the mice were i.v. injected with 0.4µg/mL PoP in 400µL 5% dextrose solution. For imaging, mice were placed on a heating pad and anesthetized with Isoflurane. The injection was left to diffuse within the tumor for 1 hour before fluorescence imaging with 660nm LED through the 740nm EMCCD filter. A treatment light at 660nm was applied to the injection site for 5 minutes, after which fluorescence images were taken again using the same parameters.

### 4.5. Preparation of long-circulating Dox in PoP Liposomes

The details of our PoP Dox liposome formulation have been described in [40, 45, 54]. PoP-liposomes incorporated PoP photosensitizer in a recent study for the development of improved peptide-based cancer vaccines, underscoring the versatility of the drug delivery platform [43]. Briefly, PoP-liposomes were synthesized from pyro-lipid through esterification of pyro with lyso-C16-PC, using 1-Ethyl-3-(3-dimethylaminopropyl) carbodiimide (EDC) and 4-dimethylaminopyridine (DMAP) in chloroform. The liposomes were formed by dispersing Porphyrin-lipid, PEGylated-lipid, cholesterol, and distearoylphosphatidylcholine in chloroform, followed by solvent evaporation. A 20 mg/mL lipid solution was extruded through a high-pressure lipid extruder with a 250 mM ammonium sulfate solution using polycarbonate membranes of 0.2, 0.1, and 0.08 µm pore size, sequentially stacked and passed through the extruder 10 times. Free ammonium sulfate was removed by overnight dialysis in a 10% sucrose solution with 10 mM HEPES at pH 7. Dox was loaded by incubating the liposomes at 60°C for 1 hour, achieving a loading efficacy of over 90% as confirmed by G-75 column tests. The self-assembly status and elution position of PoP-liposomes were tracked using 420 nm excitation and 670 nm emission, while Dox was detected using 480 nm excitation and 590 nm emission in a fluorescence plate reader (TECAN Safire).

### 4.6. Dox release quantification and PoP photobleaching by laparoscopic SFDI in mouse model

A recently sacrificed BALB/c mouse was acquired for simulated in vivo measurements. The mouse was placed on an imaging platform with the shaved injection site centered within the 3.2×3.2cm FOV of the laparoscope. SFDI measurements at the 490nm excitation and 590nm emission wavelengths of Doxorubicin were performed prior to injection. The mouse was injected subcutaneously with 50 µL of a lightly scattering intralipid medium (µ _s_’ = 5cm^-1^at 490nm) containing 23.1µg/mL PoP-Liposomes, and SFDI measurements were performed again post injection. A treatment light at 657nm at a fluence rate of 330mW/cm^2^ was directed onto a 1cm area centered on the injection site, and fluorescence measurements were acquired every 1 minute to assess drug release dynamics. PoP fluorescence images were acquired immediately before and after applying treatment light to assess photobleaching.

### 4.7. Attenuation-Compensated Fluorescence Analysis

By using SFDI in fluorescence imaging mode, photosensitizer (PS) fluorescence can allow quantification of PS concentration by accurately compensating for light attenuation at both excitation and emission wavelengths. The quantification of PoP fluorescence concentration was performed using the Gardner model [55], which corrects the raw fluorescence signal by accounting for optical absorption (µ_a_) and scattering (µ_s_’) losses at both excitation and emission wavelengths. In this model, the fluorescence correction factor, X_1D_(ex, em), is determined using the quantified optical parameters from SFDI measurements, where X_1D_(ex, em), represents the effective path length during the penetration of excitation light and the escape of emitted fluorescence from the tissue. The corrected fluorescence was then calculated as F_corr_= F_raw_/X_1D_, where F_raw_ is the measured raw signal, X_1D_ is the correction factor, and F_corr_ is the fluorescence corrected for optical absorption, scattering, and light propagation. Using the calibration factor, the corrected fluorescence was translated into PoP concentrations. Fluorescence imaging can also be used for monitoring PDT response because PS fluorescence changes during PDT, and these changes may be indicative of PDT response.

To quantitatively compare tumor regions with surrounding normal tissue, image analysis was performed using a hand-drawing tool function (*imfreehand/impoly*) in MATLAB (MathWorks, Inc., Natick, MA, USA). Regions of interest (ROIs) for both tumor and peripheral tissue were selected based on reflectance maps at 590, 625, 660, and 740 nm. Statistical metrics, including mean and standard deviation, for each ROI are summarized in a bar plot. The analysis revealed contrasts between tumor and peripheral tissue, with tumor ROIs exhibiting higher mean absorption parameters and lower mean scattering. laparoscopic SFDI data was processed similarly, with ROIs pertinent to the injected drug area being selected for analysis at each fluorescence acquisition. The mean value of this area for each minute was used to determine Dox release kinetics, and images taken at the PoP fluorescence wavelength before and after were compared to assess Porphyrin photobleaching. Data is summarized with Dox release leveling off after sufficient light dosage, and Porphyrin fluorescence decreasing due to photobleaching by drug activation light.

## 5. Conclusions

This study demonstrates the results of a wide-field laparoscopic SFDI system for real-time intraoperative monitoring of Dox-PoP. The system accurately measures PoP photobleaching and Dox release kinetics by enabling quantitative fluorescence imaging and overcoming tissue absorption and scattering limitations. The observed PDT-mediated PoP photobleaching and light-triggered drug activation confirm the laparoscopic SFDI system’s capability to enhance targeted tumor treatment while minimizing off-target effects. Compared to traditional flexible endoscopic imaging, this approach offers larger FOV and precise optical property quantification, making it a promising tool for minimally invasive photodynamic therapy and chemotherapeutic drug delivery. These findings highlight the potential of laparoscopic SFDI to improve CPT efficacy, ultimately advancing real-time monitoring of cancer treatment. Future studies should investigate its application in clinical settings and explore its integration with other imaging modalities for comprehensive tumor characterization.

## Acknowledgments

The authors acknowledge the funding support from NIH/NCI R01 (5R01CA243164-06).

## Institutional Review Board Statement

“Not applicable.”

## Informed Consent Statement

“Not applicable.”

## Data Availability Statement

The data that support the findings of this study are available from the corresponding author upon reasonable request.

